# The influence of cell morphology on the dynamics and stability of model bacterial communities

**DOI:** 10.64898/2026.02.25.707998

**Authors:** Ing Xhu Lim, Faisal Halabeya, Joshua N Milstein

**Affiliations:** Department of Chemical and Physical Sciences, University of Toronto Mississauga, Mississauga, ON, CAN; Department of Physics, University of Toronto, Toronto, ON, CAN

## Abstract

Understanding how structures arise in microbial communities remains an outstanding challenge in bacterial ecology. When replicating within dense and confined environments, bacterial populations may collectively self-organize as a result of mechanical interactions. Model dual-populations of bacteria, colonizing open-ended microchannels, were found to either quickly fixate to a single population or segregate into long-lived, coexisting populations. This latter quasi-stability results from the alignment of bacteria into lanes, forcing cells to grow toward the open ends of the channel, with inter-lane invasion events driving the boundary between populations. Here we apply agent-based (AB) simulations to explore the boundary dynamics between dual-bacterial populations of varying morphology and division rate. We find that the simulated boundary dynamics are well described by a simple drift-diffusion model, which enables us to estimate the mean time to fixation within competitive scenarios where the fixation time is difficult to access by AB simulations. Coccus cells display a competitive advantage over bacillus cells as they can more effectively invade and ultimately fixate, while coexisting populations of bacillus cells are ‘effectively’ stable. And while faster-dividing cells should have a selective advantage, their morphologydetermines if this advantage drives fixation or acts as a defensive strategy to maintain their population. These findings highlight the critical role that cell morphology and mechanics play in shaping bacterial communities.

**Author summary:** Bacteria are often found in multi-species communities, many of which are critical to human health, the environment, and industry. Within these communities, a competition for resources shapes the overall population structure and resulting ecology. Model systems can provide a test bed for understanding the interaction networks of these more complex communities. Here we show how mechanical forces between cells and their environment can be critical to the stability (or instability) of a bacterial community. Using simulations of dual species competition within two-dimensional microchannels, along with mathematical modeling, we find that cell morphology can be a determining factor in the stability of a multi-species community.

## Introduction

Bacteria commonly exist within communities in nature where they compete for limited space and resources. These communities play an essential role in supporting a diversity of critical ecosystems, from regulating metabolism and protecting against pathogens within the human gut, to cycling carbon and nitrogen within Earth’s atmosphere [1, 2]. Even initially well mixed populations of bacteria will tend to segregate and spatially organize as cells grow and divide. This resulting spatial structure may be crucial for cooperation within species [3], the sharing of public goods such as digestive enzymes or metabolites [4], or as a resistance strategy toward antibiotic compounds [5, 6]. While the availability of nutrients and the chemical environment both affect the spatial organization of bacterial communities, within dense systems, spatial structure may also emerge spontaneously through mechanical interactions with neighbouring cells and their confines [7, 8]. This emergent behaviour can critically impact the evolution and ecology of these populations.

Synthetic microfluidic systems serve as a controlled test-bed for studying self-organized spatial structuring within bacterial communities [9]. We previously reported on the rapid emergence of quasi-stable, coexisting communities within a model competitive system composed of two distinct *E. coli* strains confined to open-ended, monolayer microchambers [10]. Cells seeded within these channels would either rapidly fixate to a homogeneous population of one cell type (typically, in the time it takes for the populations to fill the channel, i.e., ∼ 6 − 8 hrs), or would organize into distinct, quasi-stable populations. Once established, these segregated populations would coexist for extended times despite significant heterogeneity between strains in terms of growth advantage or cell morphology (> 24 hrs).

These observations are largely explained by noting that, once the channel fills, the cells primarily self-align into lanes that grow toward the open ends of the channel. This arrangement is only altered by the rare cell invading and fixating in a neighboring lane. We previously showed that successful invasions are, in fact, exceedingly rare for all attempts except those that occur in the middle of a lane, away from the open ends of the channel [11]. Most invading cells are quickly pushed out of the ends of the channel, which is why heterogeneous communities with a significant difference in growth rate between populations can coexist for extended periods.

Employing both agent-based (AB) simulations and mathematical modeling, we now explore the stability of these segregated communities by studying the dynamics of the boundary between the populations. In particular, we infer the mean first-passage time of the boundary to the channel edges, which indicates the average time it takes for the community to eventually fixate to a single, homogeneous population. The boundary dynamics are found to be well modeled by a one-dimensional drift-diffusion equation whose parameters are influenced by the morphology and division rate of the bacteria.

Here we focus on competition between two bacterial strains displaying morphologies ranging from bacillus (pill-shaped) to coccus (round). These morphological differences can impart a selective advantage to the more coccus cells, such that the coccus population fixates within a finite period, whereas competition between increasingly extended bacillus morphologies leads to mean fixation times that appear to diverge. And while increased cell division can sometimes drive fixation, for well-aligned bacillus populations, it can at most stabilize the population by protecting it from invasion. Despite the simplicity of the system under study, our results highlight the role that bacterial morphology can play in the stability or instability of a bacterial community.

### Stability and boundary dynamics

We first employ agent-based (AB) simulations to model bacterial organization within dense communities that grow in two-dimensional, open-ended microchannels. AB simulations are a common tool for modelling microbial communities, and they are widely used to study population dynamics, community interactions, and emergent phenomena. [12, 13] The AB simulations used here have been described in detail within a previous publication where they were shown to capture much of the behaviour observed in actual experiments [10]. While this previous work focused on understanding the relatively rapid dynamics of the populations either fixating to a single-species or establishing segregated communities, here we focus on the longer term dynamics of’ the latter, quasi-stable communities (states we will refer to as ‘coexistence’). All simulations are initialized with one cell of each of the two competitor strains randomly seeded within a microchannel (Fig. 1A) where they are allowed to grow and divide (Fig. 1B). We then ignore all populations that fixate to a single strain within 24 hrs (Fig. 1C). The remaining systems will all have reached coexistence with the vast majority segregated into two populations separated by a boundary that runs along the long axis of the channel (Fig. 1D). Bacillus cells will tend to organize into aligned lanes that span the channel, while more coccus-like cells will display a less ordered arrangement. Occasionally, we observe segregation into 3 distinct populations (< 10 % of simulations) or into 4 or more populations (less than < 1 % of simulations), but we will neglect these configurations in our initial analysis.

**Fig 1.**
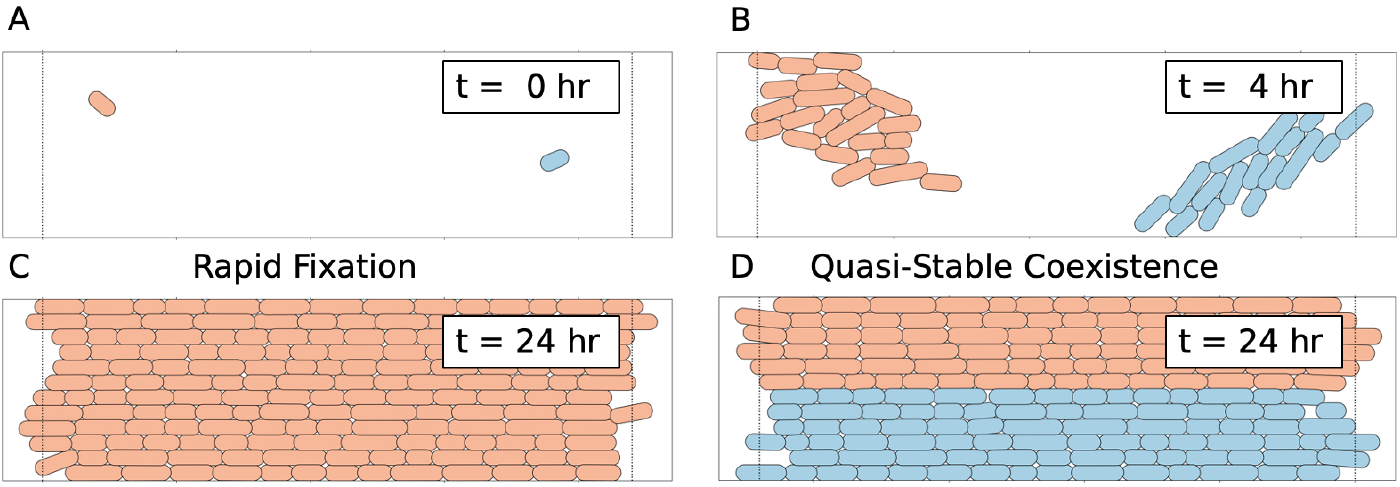
A) Cells are sparsely seeded within each channel with one cell of each strain randomly positioned and orientated. The dashed lines represent the open ends of the channel; once a cell crosses this line, it is removed from the simulation. B) Cells then spread out within the chamber due to multiple rounds of cell division. This will eventually lead to either C) fixation of a single strain or D) a quasi-stable, coexisting state where different populations, each composed of a single strain organized into lanes, segregate within the chamber. Cell division now primarily drives growth toward the open ends of the chamber.

We simulated competition between populations of cells with morphologies ranging from bacillus (pill-shaped) to coccus (round). The cells were modelled as rectangular segments of finite length *l* and 1 µm width with half-circular end caps 1 µm in radius (see S1 File). By increasing *l*, we move from coccus to increasingly elongated bacillus cells. As an example, Fig. 2A illustrates competition between perfectly coccus cells and bacillus cells 2.5 µm long on average. Figures 2B,C display the rapid fixation of the systems to either a pure bacillus or coccus population, respectively, while Fig. 2D displays the comparatively slower dynamics of the two populations within the coexistence state. Throughout, we will employ the fractional area, which is the total area of a population divided by the area of the microchannel, as a convenient measure of the population abundance. Note, however, that due to edge effects at the channel boundaries, this number can fluctuate above 1.

**Fig 2.**
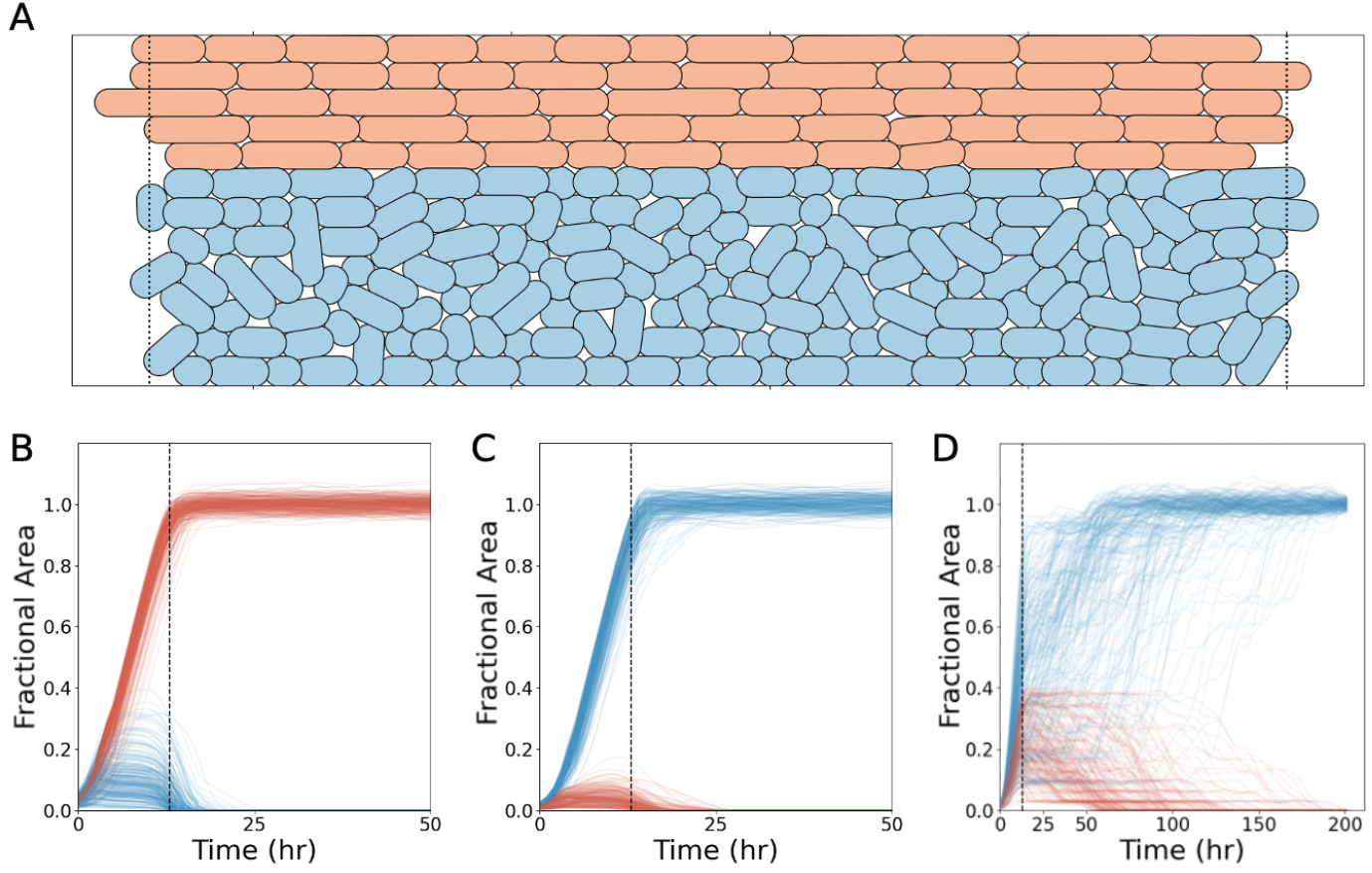
A) During coexistence, the boundary between segregated populations evolves through repeated invasion attempts by cells at the boundary. Coccus cells (blue), with less of a propensity to self-align, are more likely to push into the domain of bacillus cells (red). We can separate the competitive outcomes into B) and C) fast (< 24 hrs) and D) slow (> 24 hrs) fixation events. Fast fixations occur when one population out-competes the other in an effort to fill the chamber, while slow fixations result from the boundary dynamics between segregated populations. For this example, the bacillus cells never out-compete the coccus cells once coexistence is reached.

Here we are primarily interested in the dynamical process of fixation from the coexistence state, which is driven by the motion of the boundary separating the two populations. To illustrate this, Fig. 3A displays a kymograph of the relative position of the boundary over time showing a series of discrete jumps that correspond to successful invasions into one or more lanes of the neighboring bacillus population. These discrete jumps are also observable in a plot of the fractional area versus time. The takeover of the bacillus cells by the coccus cells occurs in a predictable way as illustrated in Fig. 3B. First, coccus cells invade into the neighbouring lane of bacillus cells with successful invasions almost always occurring near the centre of the chamber. Then, as the invading cells divide, their progeny slowly push the bacillus cells towards the open ends of the chamber until all bacillus cells in that lane have been expelled. These steps repeat, interspersed with many unsuccessful invasion attempts, until the simulation ends or the coccus cells take over the entire chamber.

**Fig 3.**
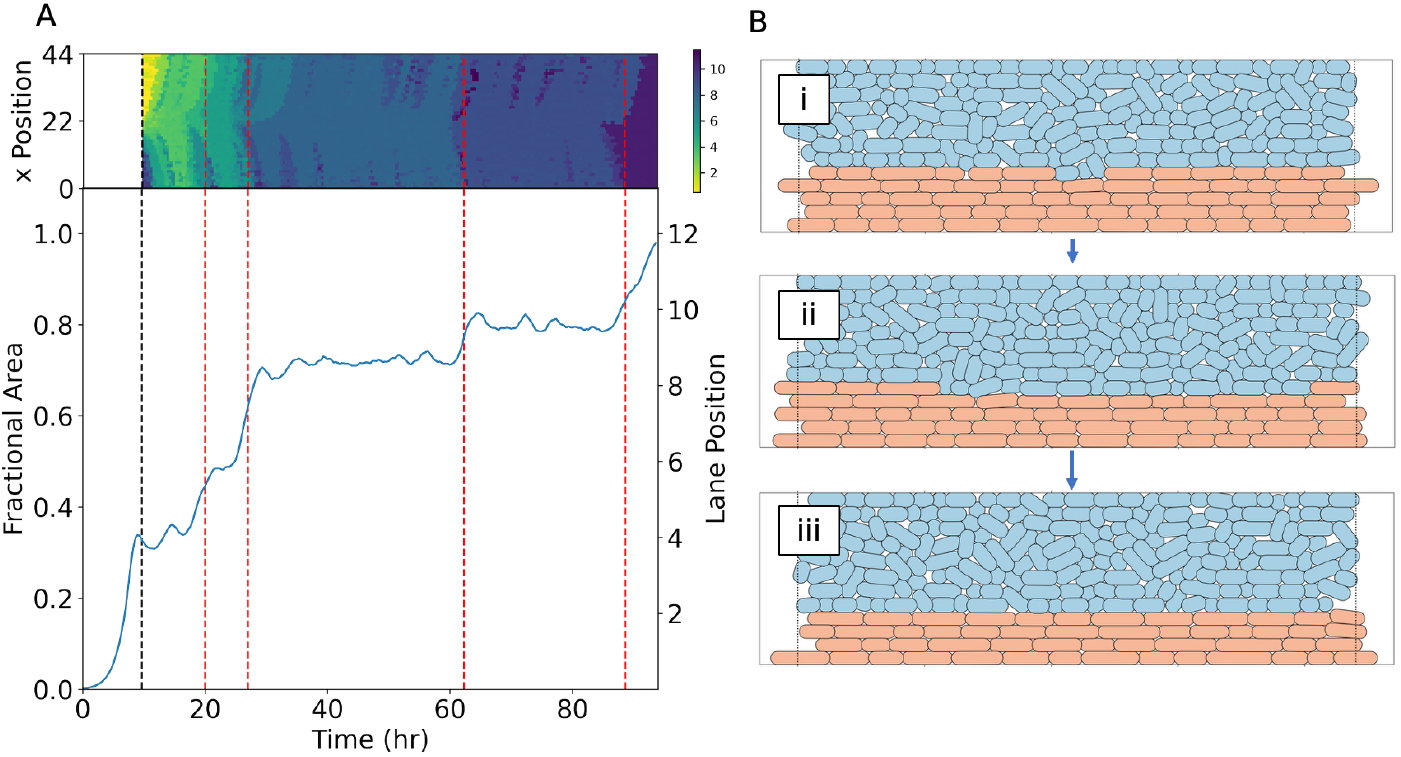
A) Boundary dynamics from simulations of bacillus (2.5 µm) versus coccus (1 µm) cells. Top panel: Kymograph of the boundary dynamics as a function of time. The colours represent the spatial location of the boundary discretized into lanes within a channel supporting up to 12 lanes. Bottom panel: Fractional area with change-point analysis identifying successful invasions (red-dashed vertical lines) showing the boundary move in discrete jumps. The black-dashed vertical line indicates the chamber fill time. B) Boundary dynamics are driven by invasions. Illustrated here: i) coccus cells invade a lane of bacillus cells near the chamber centre, ii) cell division by the invaders then push the competing bacillus cells toward the edges of the chamber, and iii) the lane fixates to coccus cells shifting the location of the boundary downward. The dashed vertical lines indicate the chamber boundaries.

Since the timescale to fixation of a single lane, post invasion, is typically much faster than the translational motion of the boundary (Fig. 3B), we may ignore the intra-lane fixation dynamics and approximate the boundary as moving in discrete jumps between lanes. Within this approximation, as we will show, the translational dynamics of the boundary may be modelled as simple drift-diffusive motion. Employing AB simulations to extract both the diffusion constants and drift velocities, for boundaries between cells of varying morphology, we first show that the model predictions agree well with AB simulations of systems that reach fixation from coexistence within about a week (corresponding, roughly, with the actual run-time of the simulations). However, our analytical model also enables us to predict the time to fixation for competitive systems when the fixation time becomes extremely long, making it difficult to assess through AB simulations alone.

### Drift and diffusion of the population boundary

The translational motion of the population boundary can be described by the Fokker-Planck equation, which characterizes the spatial and temporal evolution of the probability density function *P*(*x, t*) for the boundary:

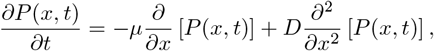

where *t* is time, *x* is a spatial coordinate indicating the location of the boundary between the channel walls, *µ* represents the drift velocity, and *D* is a diffusion constant. For the moment, we assume that both *µ* and *D* are constants. From repeated runs of the AB simulations, estimates of these parameters can be inferred from the simulated data using the following relationships:

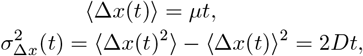

where Δ*x* is the relative displacement from the boundary’s initial position:

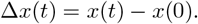

### Effects of relative morphology and division time

We considered AB simulations of two bacterial populations with one population held fixed at an initial cell length of 2.5 µm and the other varied from an initial length of 1.0 µm (coccus) to 2.5 µm, as well as between two coccus populations (1.0 µm vs. 1.0 µm). Cells grew to double their initial length and divided at equal rates of one division every 60 ± 6 minutes. Gaussian noise (10%) was introduced in the division rate to prevent synchronization of cell size and growth stage (see S1 File). Figures 4A-B display the observed dynamics of the relative displacement of the boundary Δ*x*(*t*) for the first 40 hrs post-coexistence for the cases: 2.5 µm vs. 1.0 µm, and 1.0 µm vs. 1.0 µm (see S1 Fig). Once again, the relative area is used as a measure of the boundary coordinate *x*(*t*). A linear fit to the mean of the trajectories yields the drift velocity *µ*. Similarly, the inserts show the evolution of the variance 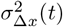 for the first 40 hrs post-coexistence, which can be fit to a line to extract the diffusion constant *D*.

**Fig 4.**
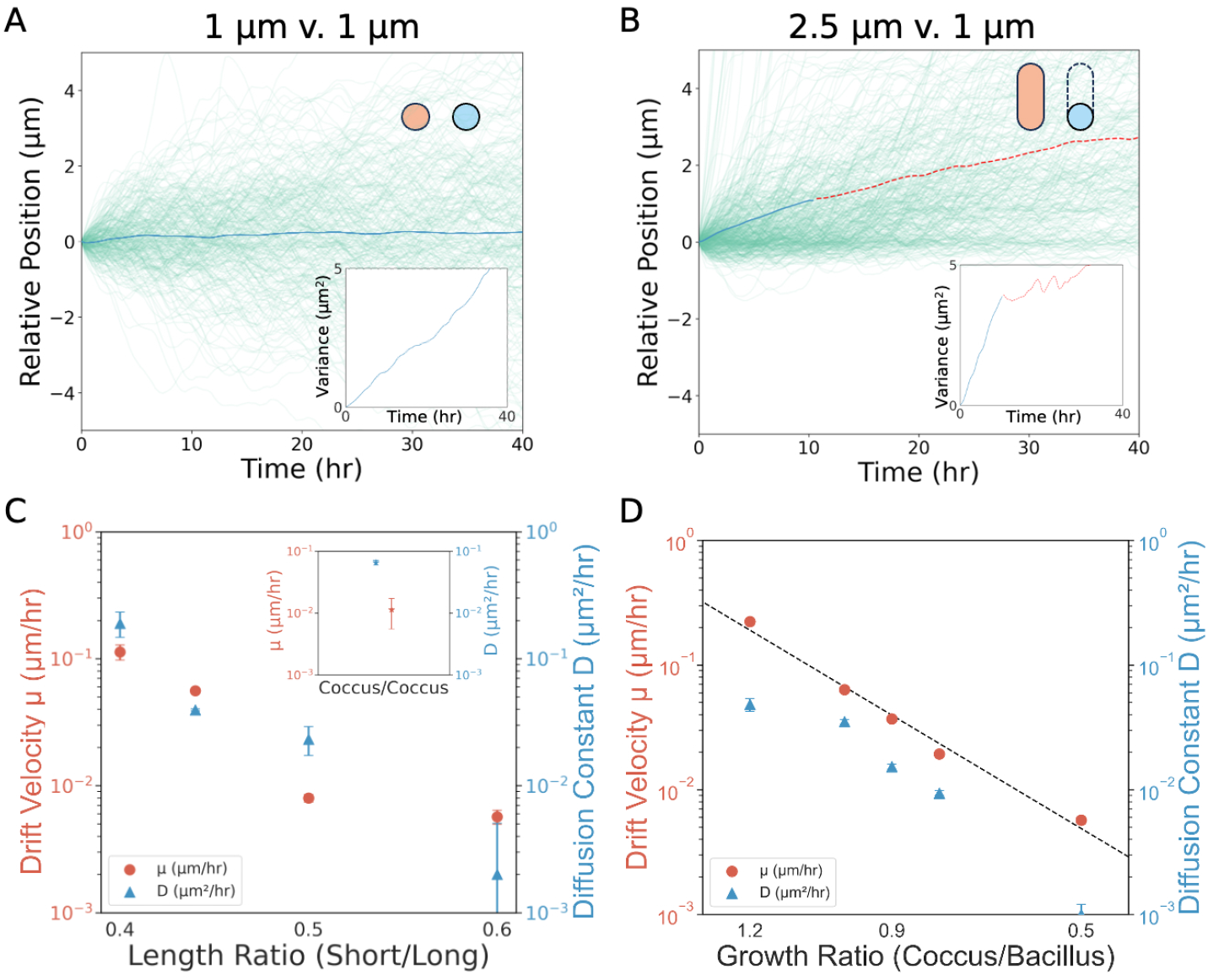
Relative displacement of the population boundary as a function of time for competition between populations with the following morphologies (defined by the initial cell lengths): A) 1.0 µm vs. 1.0 µm and B) 2.5 µm vs. 1.0 µm. Shown are the simulated trajectories of the boundary (green lines) as well as the mean of the trajectories (blue line); the data is no longer fitted due to many simulations terminating, thus reducing sampling (red dashed line). Insets display the corresponding variance over time. Linear regressions quantify: (i) drift velocity (mean slope) and (ii) diffusion coefficient (variance slope). **(C)** Drift velocities (red) and diffusion coefficients (blue) as a function of morphology relative to the longest bacillus cell (length/2.5 µm). Data points shown are for size ratios of 0.6, 0.5, 0.44, and 0.4. Inset: Same for 1.0 µm vs. 1.0 µm (coccus/coccus) competition. **(D)** Drift velocities (red) and diffusion coefficients (blue) as a function of relative growth rate for 2.5 µm vs. 1.0 µm competition (size ratios of 0.5, 0.8, 0.9, 1.0, and 1.2), with the growth rate of coccus cells held fixed. Each data point in C and D was obtained from 1000 simulations, as were the graphs in A and B.

Figure 4C summarizes the drift velocities (*µ*) and diffusion constants (*D*) extracted from simulations as a function of relative morphology (i.e., with one population held at 2.5 µm (long) and the other varied at shorter lengths (short)), but only for those cases where we observed a significant number of lane changes, thus allowing us to reliably extract the diffusive parameters. Morphology is seen to provide a strong competitive advantage, with the drift velocity—which largely determines the dynamics—increasing by orders of magnitude as bacillus cells compete with increasingly coccus cells. Under these conditions, with the drift always promoting invasion of the bacillus population, the more coccus cells dominantly fixate. Conversely, any dynamics between comparable bacilli populations is exceedingly slow, effectively jamming the two populations into a coexisting state. For competition between dual coccus populations, as expected, the dynamics of the boundary are largely diffusive (insert Fig. 4C).

We next considered the effects of varying the relative division time between populations. We analyzed AB simulations of competition between elongated bacillus and coccus cells (2.5 µm vs. 1.0 µm) keeping the mean division time of the coccus populations constant (60 ± 6 minutes) while ramping up the division time of the bacillus cells and maintaining Gaussian noise of ±10% about the mean. In Fig. 4, we display the resulting diffusion parameters as we decreased the relative division time (*i*.*e*., growth rate ratio (coccus/bacillus)) by a factor of 2 (see S2 Fig).

While the drift velocity of the boundary decreased by orders of magnitude as the division time of the bacillus population doubled, the drift remained into the bacillus population, meaning that the coccus cells were still the population to dominantly fixate. A faster division time, while not contributing to an increase in self fixation, instead protected the bacillus cells for longer from colonization and eventual fixation by the coccus population. Coccus cells, attempting to colonize the bacillus population, are instead rapidly pushed to the boundary and ejected from the population before they are able to take over a new lane. In our simulations, we observed an exponential dependence of the drift of the coccus population as a function of division time (exponent *α* = −7.9). While the value of the exponent is likely influenced by details of our ABM simulations, the qualitative observation of an exponential dependence on the division dynamics illustrates how the rate of cell division can significantly alter population dynamics.

### Mean first-passage times

For one-dimensional diffusion, where both the drift *µ* and diffusion *D* are constant, we can derive a second-order differential equation for the mean first-passage time *τ*(*q*) [14],

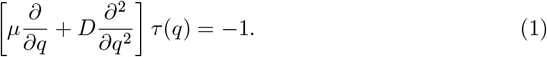

where 0 ≤ *q* ≤ *W* specifies the starting point between the two walls of the microchannel. For absorbing boundary conditions such that *τ* (0) = *τ*(*W*) = 0, this can be solved to yield the rescaled mean first-passage time (MFPT):

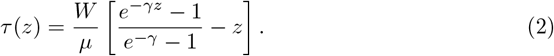

Here *γ* = *µW/D* and we scale the starting position of the boundary to the chamber width *z* = *q/W*. In the limit of zero drift (*U* → 0) this becomes

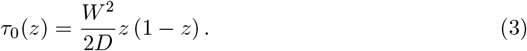

Employing the diffusive parameters found from our AB simulations, we may use Eq. 2 to predict the MFPTs for the various competitive scenarios shown in Fig. 4.

To validate the derived MFPT model, theoretical predictions were compared with the empirical data obtained from simulations for the 2.5 µm vs. 1.0 µm cells. For this scenario, all populations fully fixated from coexistence within the maximum simulated time of 200 hrs. This happens to roughly correspond to the simulation run time, or a little over a week on our computing cluster. A comparison of the fixation times displayed in simulations and MFPT predictions, for possible initial positions of the population boundary, are shown (inset Fig. 5A).

**Fig 5.**
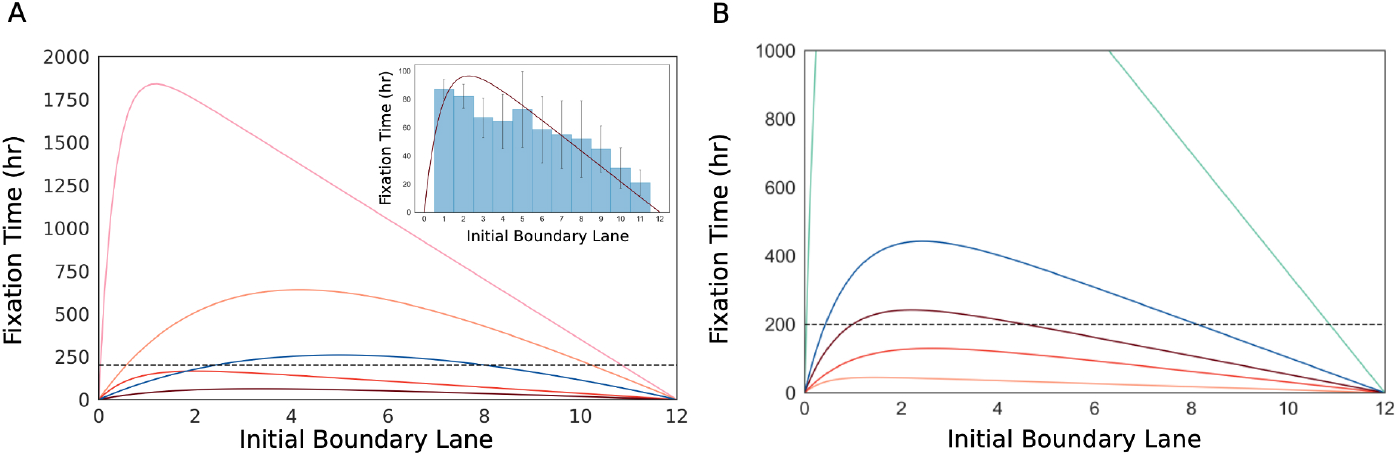
Theoretical predictions of fixation times for population boundaries that initialize at varying positions between the channel walls. This is shown A) as a function of cell morphology ratios (top to bottom: 0.6, 0.5, 0.44, 0.4) and B) as a function of relative division time ratios (top to bottom: 0.5, 0.8, 0.9, 1.0, 1.2). A simulation cutoff time of 200 hrs is indicated (black horizontal dashed line). The insert in (A) displays a histogram of the fixation times for 2.5 µm vs 1.0 µm competition at a growth rate ratio of 1. Bars represent the average fixation times at each initial boundary position, with the curve indicating the predicted MFPTs.

Most of the simulations, at less distinct relative morphologies, did not fixate a sufficient number of times during the simulation to extract the full MFPT distribution. For these instances, using the drift-diffusion parameters from the previous section, we simply employ our MFPT model to predict the fixation times. Figure 5A shows the predicted fixation times as a function of initial boundary position for competition between size ratios of 0.6, 0.5, 0.44, and 0.4. As we vary the length of the initially coccus population toward that of the 2.5 µm bacillus competitor cells, the predicted fixation times rapidly increase, leading to an increasingly stable coexistence of the two populations. Likewise, the predicted fixation times for competition between 2.5 µm bacillus vs. 1.0 µm coccus cells, when varying the relative division time between populations, are shown in Fig. 5B. As the division time of the bacillus population is decreased, the time to fixation of the coccus population, from coexistence, is increasingly extended.

## Conclusions

In this study, employing agent-based simulations and mathematical modelling, we investigated the role of bacterial morphology in determining the stability of spatially structured microbial communities confined within a 2D monolayer. Within these dual-population systems, cells tend to spatially order into distinct lanes of a given cell type with competition driven by successful inter-lane invasions.

We find that rounder, more coccus-like cells consistently display an advantage over elongated, pill-shaped bacillus cells due to an increased affinity for invading neighbouring lanes, which more effectively drives the system toward fixation. This competitive advantage can be mitigated by increasing the growth rate of the bacillus cells. However, doing so does not promote fixation of the bacillus population; rather, it only protects the population from invasion and eventual fixation by the more coccus competitor strain.

A simple one-dimensional drift-diffusion model for the boundary between populations effectively captures the long-term fixation dynamics. Here we employ this model to predict the fixation time for systems whose boundary dynamics become exceedingly slow, such that the populations form a jammed or quasi-stable equilibrium, which is the case for competition between similarly elongated bacillus populations. These slow dynamics are difficult to access through AB simulations alone.

One could argue that more sophisticated algorithms than those we employed might provide access to longer time scales. Still, we have shown that both morphology and division dynamics can greatly stabilize these mixed communities. The question remains, however, as to whether these competitive systems are ever truly stable such that the fixation time approaches infinity. Likewise, here we have relied on our phenomenological AB simulations to infer dynamical parameters. More realistic AB simulations, that incorporate the microscopic interactions between cells and the environment, could provide further insight into these systems. And one should, ideally, be able to extract these dynamical parameters from a microscopic model alone.

Collectively, our findings highlight the influence that cell morphology has on the stability of competing bacterial communities. Morphological traits, through purely mechanical interactions, can strongly influence the long-term competitive outcomes and ecological trajectories of bacterial populations. Morphology can give rise to dynamic and spontaneous structure within bacterial communities, which can affect competitive dynamics in unexpected ways. Within these structures, seemingly obvious advantages—such as being able to replicate faster—may not always lead to the dominance of a population. Similarly, phenotypic traits such as cell morphology, in certain settings, may play a role in eradicating persistent populations resistant to other forms of treatment. These findings deepen our understanding of microbial ecology, provide valuable insight into how microbial communities form collective structure, and may lead to more predictive tools for exploring and managing bacterial community dynamics across natural and engineered ecosystems.

## Supporting information

**S1 File. Agent Based Models**. A brief description of the agent based models employed.

**S1 Fig. Relative position and variance for extended morphologies**. Relative displacement of the population boundary as a function of time for competition between populations with the following morphologies (defined by the initial cell lengths): A) 2.5 µm vs. 2.5 µm and B) 2.5 µm vs. 2.0 µm. C) 2.5 µm vs. 1.25 µm. D) 2.5 µm vs. 1.75 µm. E) 2.5 µm vs. 1.5 µm. F) 2.5 µm vs. 1.1 µm. Shown are the simulated trajectories of the boundary (green lines) as well as the mean of the trajectories (blue line); the data is no longer fitted due to many simulations terminating, thus reducing sampling (red dashed line).

**S2 Fig. Relative position and variance for extended growth ratios**. Relative displacement of the population boundary as a function of time for competition between populations with the following growth rate ratio (defined by the ratio of coccus (1.0 µm)/bacillus (2.5 µm)): A) 0.9 and B) 1.2. C) 0.8. D) 0.5. Shown are the simulated trajectories of the boundary (green lines) as well as the mean of the trajectories (blue line). Each are taken from 1000 simulations, however, as the growth rate changes the percentage of seeded simulations that reach the quasi-stable, coexistence phase decrease.

## Acknowledgments

We would like to thank John Whitfield for his contributions during the early stages of this research and Jeremy Rothschild for developing the AB simulations we employed.

## Notes

https://doi.org/10.5683/SP3/VEKDFA

### Competing Interest Statement

The authors have declared no competing interest.

