## Supplemental File 1 for "The influence of cell morphology on the dynamics and stability of model bacterial communities"

**Agent Based Model**

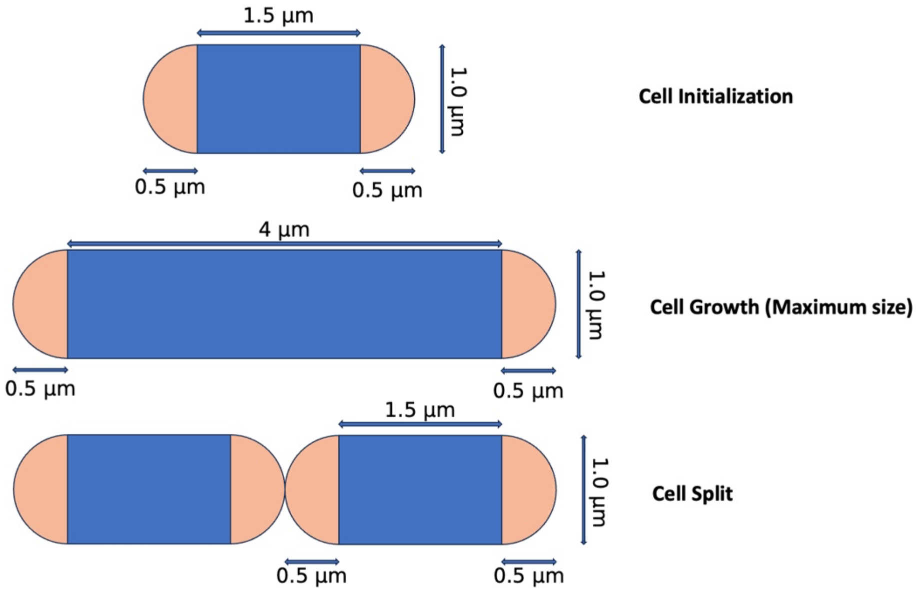


**Figure S1.1:** Illustration of cell growth and division within the ABM.

The Agent based Model (ABM) is a continuation of the work done by Ma et al. (PNAS, 2024), utilizing much of the same configuration. In a virtual environment, the model aims to simulate cell-cell mechanical interactions by allowing each cell to exist through a governing set of rules that determine their growth, division, and interactions with their environment and neighbours. By including mechanical pushing among neighbouring cells, the model captures the cooperative and competitive behaviours that shape bacterial colonies' spatial distribution and growth trajectories. This approach is used again within our experiments to understand the spacial dynamics of competing cells populations.

The ABM simulated a 2D rectangular environment with dimensions 44 um x12 um where cells are grown within a monolayer. The bacterial cells are modeled as agents of spherocylindrical shape with maximal length, and growth rate. Cells grow by exponentially increasing the length of the rectangular portion of the Spherocylinder until the total length of the cell matches the desired maximum length--by then the cell is split into two equal sized new Spherocylinders. Physical contact between the cells happens when cells overlap with each other causing a repulsion Hertzian force within a confined, damped environment. The mechanical details implemented in our ABM are provided in Ma et al. (PNAS, 2024). Each cell persists in the simulation until its centre exits the microchannel boundaries through either of the open ends. At this point, the cell is deemed to have exited the channel and is consequently eliminated from the simulation.

In previous work, the maximal length was drawn from a uniform distribution of final lengths. Here, we instead fix the final length. However, this resulted in synchronized simulations that gave rise to artifacts such as oscillations in the data. Instead, to make the cells asynchronous, we draw the growth rates from a gaussian distribution with a standard deviation of 10% of the cells growth rate around the defined growth rate.
