## Supplementary figures and images for "The influence of cell morphology on the dynamics and stability of model bacterial communities"

### Supplemental Figure 1

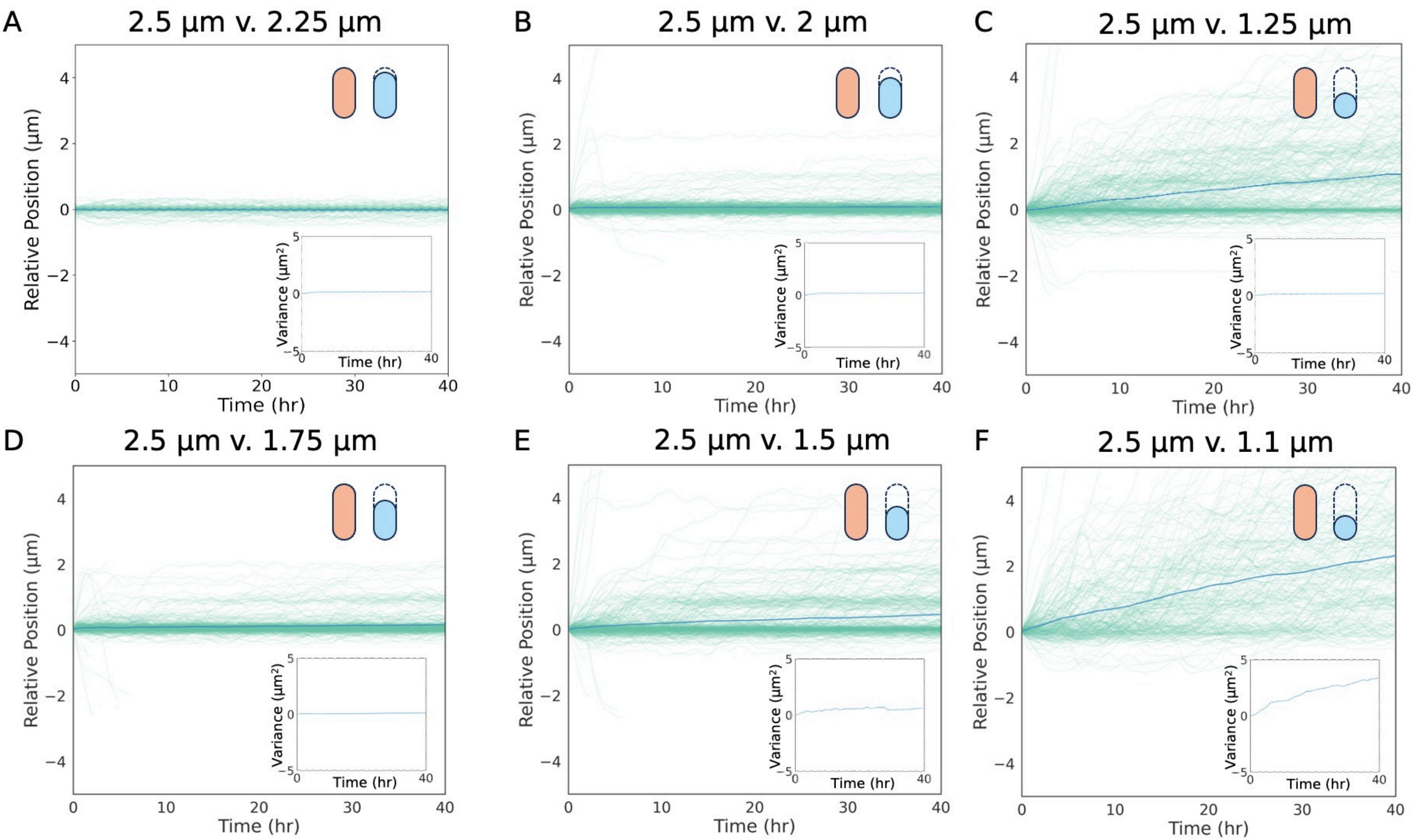

### Supplemental Figure 2

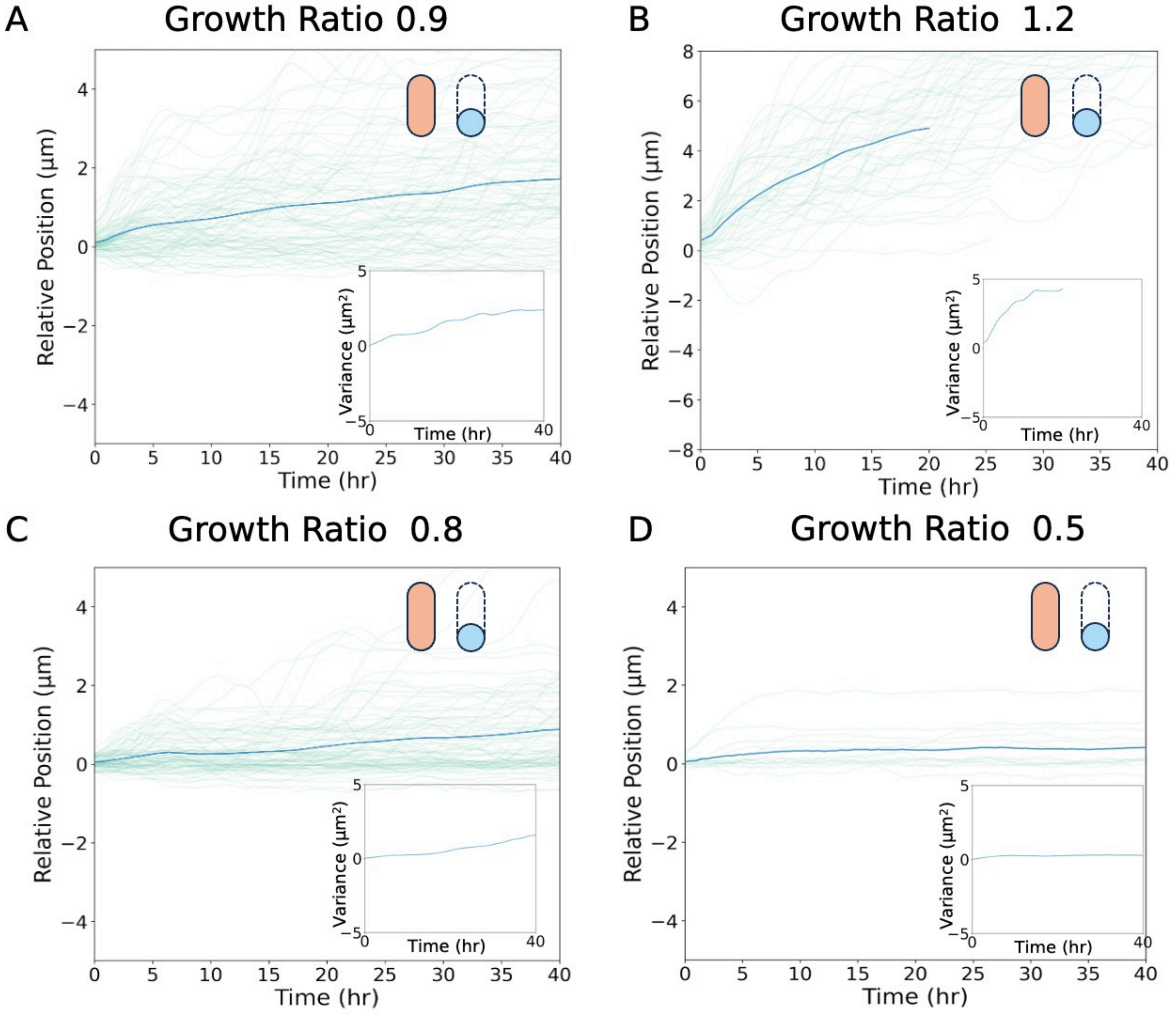
